# Association of maternal prenatal psychological stressors and distress with maternal and early infant faecal bacterial profile

**DOI:** 10.1101/713602

**Authors:** Petrus J.W. Naudé, Shantelle Claassen-Weitz, Sugnet Gardner-Lubbe, Gerrit Botha, Mamadou Kaba, Heather J. Zar, Mark P. Nicol, Dan J. Stein

**Author notes:** Authors contributed equally to this manuscript (Petrus J.W. Naudé, Shantelle Claassen-Weitz). Corresponding author: P.J.W. Naudé, Department of Psychiatry and Mental Health and Neuroscience Institute, Faculty of Health Sciences, University of Cape Town, Cape Town, South Africa. Outpatient Building, H- Floor Research Offices, Circle Groote Schuur Drive, Groote Schuur Hospital, Observatory, 7925, Cape Town.

## Abstract

**Background:** Findings from animal studies indicate that the early gut bacteriome is a potential mechanism linking maternal prenatal stress with health trajectories in offspring. However, clinical studies are scarce and the associations of maternal psychological profiles with the early infant faecal bacteriome is unknown. This study aimed to investigate the associations of prenatal stressors and distress with early infant faecal bacterial profiles in a South African birth cohort study.

**Methods:** Associations between prenatal symptoms of depression, distress, intimate partner violence (IPV) and posttraumatic stress-disorder (PTSD) and faecal bacterial profiles were evaluated in meconium and subsequent stool specimens from 84 mothers and 101 infants at birth, and longitudinally from a subset of 69 and 36 infants at 4–12 and 20–28 weeks of age, respectively in a South African birth cohort study.

**Results:** Infants born to mothers exposed to high levels of IPV had significantly higher proportions of unclassified genera within the family Enterobacteriaceae detected at birth; higher proportions of the genus *Weissella* at 4-12 weeks; and increased proportions of genera within the family Enterobacteriaceae over time (birth to 28 weeks of life). Faecal specimens from mothers exposed to IPV had higher proportions of the family Lactobacillaceae and lower proportions of Peptostreptococcaceae at birth. Maternal psychological distress was associated with decreased proportions of the family Veillonellaceae in infants at 20-28 weeks and a slower decline in Gammaproteobacteria over time. No changes in beta diversity were apparent for maternal or infant faecal bacterial profiles in relation to any of the prenatal measures for psychological adversities.

**Conclusion:** IPV during pregnancy is associated with altered bacterial profiles in infant and maternal faecal bacteria. These findings may provide insights in the involvement of the gut bacteria linking maternal psychological adversity and the maturing infant brain.

## Introduction

The prevalence of maternal prenatal stressors and distress are particularly high in rural South Africa. Previous studies reported that 30-40% of pregnant mothers have depression (1-3), 11-19% have posttraumatic stress disorder (PTSD) (4, 5) and 21-32% experienced intimate partner violence (IPV) within the past year (5, 6). Psychological stressors and distress during pregnancy may have a range of negative consequences on the development and health of the foetus and infant, including premature delivery, impaired cognitive development, poor infant growth and dysregulated behaviour of infants (7). The biological mechanisms responsible for the intergenerational transmission of the effects of maternal psychological stress during pregnancy are, however, still unclear. Studies in mice (8) and monkeys (9) have found that exposure to psychological stressors during pregnancy alters the profile of the maternal faecal microbiota, which is then transferred to the offspring. There are, however, few human studies. A Dutch human study (10) found that a higher self-reported stress during pregnancy was associated with temporal differences in infants’ faecal bacterial profiles, starting 7 days after delivery. Furthermore, infant faecal bacterial profiles from mothers reporting higher stress levels were characterized by higher relative abundances of Proteobacteria (including the Enterobacteriaceae *Escherichia, Serratia* and *Enterobacter*) and a lower relative abundances of lactic acid bacteria and Bifidobacteria (10). A recent study examined the associations of maternal prenatal anxiety, depression and stress during pregnancy with changes to the microbiome in meconium of 75 new-borns (11). Higher measures of pregnancy-related anxiety were associated with lower abundances of an unidentified genus in the family Enterobacteriaceae (11). However, the association of other types of psychological stressors and distress during pregnancy with the temporal changes in the infant faecal microbiome has not yet been studied.

The acquired faecal microbiota may contribute to child neurodevelopment and health trajectories (12-14). A delayed or altered establishment of a mature intestinal microbiota in childhood (termed microbiota immaturity) has been associated with various diseases, including diarrhoea, malnutrition, atopic conditions, inflammatory bowel disease, obesity, diabetes, neurological conditions and impaired neurodevelopment (15, 16). Studies in mice demonstrated that the gut microbiome may function as mechanistic link between maternal prenatal stress and reduction in social behavioural and neurobiological changes, i.e., neuroinflammation, in the offspring (8, 17). The early establishment of the gut bacterial environment may have long-lasting effects on health and brain functioning. In this respect, a study in germ-free mice showed that colonization of the gut directly after birth was essential for the development of the hypothalamic-pituitary-adrenocortical (HPA) axis (18). A study in humans also showed that changes in gut bacteria at one year of age were associated with with lower cognition, including visual reception and expressive language skills in infants at two years of age (16). However, evidence for the association between maternal prenatal psychological adversities with changes to the infant gut microbiome in utero, i.e., meconium, is still limited.

### Aims of the study

The primary aim of this study was to determine the association between different profiles of maternal prenatal stressors and distress (including IPV, psychological distress, depression, and PTSD) experienced by South African mothers and the development of infants’ faecal microbiome early in life. The secondary aim included the investigation of the relationship of these prenatal psychological adversities with maternal faecal bacterial profiles at delivery.

## Materials and methods

### Study participants

Mother-infant dyads were evaluated in a sub-sample from the Drakenstein Child Health Study (DCHS). The DCHS is multidisciplinary birth cohort study investigating determinants of child health over time. Further details of participant recruitment, data collection and the setting of the DCHS has been provided previously (19, 20). Pregnant women were recruited from two low socioeconomic communities, TC Newman and Mbekweni in a peri-urban area outside Cape Town, South Africa. Enrolment of pregnant women took place at 20-28 weeks of gestation at routine antenatal visits to health care facilities. Mothers provided informed, written consent for enrolment of their infants at the time of delivery and follow up until five years of age. Inclusion criteria for this sub-study were longitudinal collection of faecal specimens and availability of maternal prenatal psychological measurements. Exclusion criteria for this sub-study were maternal age < 18 years, residence outside of the Drakenstein sub-district, and intention to move out of the region within 2 years of giving birth.

This study and parent study [the Drakenstein Child Health Study (DCHS)] both received ethical approval from the Faculty of Health Sciences, Human Research Ethics Committee (HREC) of the University of Cape Town (401/2009 and 742/2013, respectively) (19).

### Measurements of maternal prenatal psychological stress and distress

Locally validated and reliable psychological measures of stress and distress were performed in pregnant mothers when enrolled at the antenatal clinic as previously described (5, 19, 20). The measures for maternal prenatal stressors and distress in this sub-study were evaluated at a gestational age of 27.4 ± 4.2 weeks by field workers in the absence of the partner/husband. On-site female fieldworkers administered a battery of self-report measures described below. Female fieldworkers were selected based on criteria previously found to affect women’s willingness to divulge intimate information, including measures of IPV exposure (21). All female fieldworkers had at least a Grade 12 certificate and had prior experience in psychiatric/psychological research. They were fluent both in English and Afrikaans or isiXhosa and were therefore able to administer questionnaires in the participants’ preferred language. Further, fieldworkers received extensive in-service training on all aspects of Good Clinical Practice (GCP) (22).

The self-reporting questionnaire 20-item (SRQ-20) (23), a WHO-endorsed measure of symptoms of psychological distress, with good face validity and reliability, has been used widely in international studies and in South African settings (24, 25). Symptoms of depression were measured by the Beck Depression Inventory (BDI-II), a widely-used and reliable measure of depressive symptoms (26, 27). The modified PTSD Symptom Scale (MPSS) is a 17-item self-report scale with excellent internal consistency, high test-retest reliability and concurrent validity to diagnose PTSD consistent with DSM-IV criteria as defined by DSM-IV (28, 29). The validity of the MPSS was established in a South African student sample with a Cronbach’s alpha of 0.92, in relation to a diagnosis by psychiatrists based on clinical interviews for PTSD, for which kappa was found to be 0.68 (30).The IPV Questionnaire used in this study was adapted from the WHO multi-country study (31) and the Women’s Health Study in Zimbabwe (32). The IPV questionnaire assess lifetime and recent (past-year) exposure to emotional, physical and sexual IPV. Each category of violence was assessed across multiple items measuring the frequency of a number of specified violent acts. Each item is scored using a frequency scale from 1 (“never”) to 4 (“many times”). Bivariable analyses were performed on scoring guidelines based on prior work in South Africa (33). Psychological assessments of IPV were previously validated and optimized in local studies of maternal mental health for their use in the South African setting (34).

### Faecal bacteria analyses

Details of the study methods related to the analyses of faecal bacteria have previously been reported (35). In short, faecal specimens were collected using sterile spatulas and faecal screw-cap containers. Study staff collected faecal specimens from mothers and infants prior to hospital discharge or during visits to clinics. All specimens collected at hospital or visits to clinics were immediately stored at -20 °C. Mothers were instructed to collect and immediately store infant faecal specimens at -20 °C in the event where faecal specimens were not passed at hospital or during scheduled visits to clinics. All home collections were transported to clinics using ice-boxes and were delivered to study staff within 24 hours of collection. Transport of faecal specimens between the study site and laboratory was performed under controlled conditions using ice boxes. Upon arrival at the laboratory, faecal specimens were stored at -80 °C until further processing. Faecal specimens included in this study were selected based on the availability of faecal specimens from mother-infant dyads at birth and at follow-up visits. Specimens were selected to include the maximum number of longitudinal sample sets at the time of study.

Nucleic acid was extracted from each of the faecal specimens using approximately 50mg of starting material as described previously (35, 36). A detailed description of the amplicon library preparation, sequencing and bio-informatics steps was published previously, including access to the raw sequence files supporting the findings of this manuscript (35). Briefly, amplicon library preparation was performed by amplifying the V4 hypervariable region of the 16S ribosomal ribonucleic acid (rRNA) gene (35). Sequencing was carried out using the Illumina® MiSeqTM platform and the MiSeq Reagent Kit v3, 600 cycles (Illumina, CA, USA) (35). Quality filtering steps of raw sequences, removal of potential contaminants, de-replication of sequences occurring more than twice, clustering of sequences into operational taxonomic units (OTUs), removal of chimeras and taxonomic assignment was performed using an in-house bio-informatics pipeline incorporating various software tools (35).

### Covariates

Data on maternal demographics (residential area, education) and health (HIV status, smoking) were obtained at enrolment. Maternal body mass index (BMI) was determined at 6–10 weeks postpartum. Data for mode of delivery, gestational age, birth weight and length, infant sex, antibiotic use and household members were obtained at the time of delivery at Paarl Hospital, where all births took place. Information on feeding practices was obtained at infant follow-up visits at 4-12, and 20-28 weeks of age (19). The covariates included in this study were chosen based on their potential effect on faecal bacterial profiles as shown previously (35). Variables that were associated with prenatal psychological measures; or bacterial taxa at phylum, class, order, family or genus-level at each of the time points under study; or diversity indices at each of the time points under study were included as covariates and referenced in the results section. Measures for other covariates, for example, maternal diet at birth, that may affect the associations investigated in this study, were not assessed in this study.

### Statistical analyses

R software version 3.1.1 (37) together with RStudio software version 0.98.50751 was used for all statistical analyses as well as graphical representations of the data. Count data (38) was transformed to compositional data by calculating the relative abundance of each OTU per specimen (39, 40). Alpha-diversities were determined using the Shannon diversity (HIZ) index (41, 42) using the vegan R package (43). Beta-diversities were computed based on the “w” metric as decribed in Koleff et al. (44) with the vegan R package and represented in a SMACOF multidimensional scaling map (45). Maternal prenatal psychological measures in this study included symptoms of depression, psychological distress, measures for IPV and PTSD. Measures for psychological distress (SRQ-20) and symptoms of depression (BDI-II) were used as continuous variables. Dichotomized measures for IPV were used and PTSD was categorized according to “trauma-exposed PTSD”, “suspected PTSD” and “no exposure to PTSD”. A multivariate approach was followed, testing simultaneously which covariates significantly influenced the microbiome composition by performing PERMANOVA (46) on the Bray-Curtis dissimilarity matrix (calculated using the [vegdist] function from the R package vegan) (42, 47-49). For those covariates identified as significant, generalized linear models (GLMs) were used to evaluate the association of prenatal psychological measures with each taxon and alpha-diversity at each of the time points under study (35). Similarly, generalized linear mixed models (GLMMs) were used to investigate covariate effects across time points using a subset of 36 infants with complete longitudinal data sets. Hypothesis testing was performed at a 5% significance level and since the GLM and GLMM models were fitted for one taxon at a time, multiple testing necessitated controlling the false discovery rate with the Benjamini-Hochberg correction (50).

## Results

### Study characteristics

Faecal specimens of 90 mothers, 107 infants at birth, 72 infants at 4–12 weeks and 36 infants at 20–28 weeks were collected and successfully sequenced (35). Maternal prenatal psychological measures were not collected from six of the mothers. These mothers and their infants (six infants at birth and three infants at 4–12 weeks) were excluded from this study. Associations between prenatal distress measures and infant faecal bacterial profiles were therefore investigated from 101 infant meconium specimens, 69 faecal specimens collected at 4-12 weeks and 36 faecal specimens collected at 20-28 weeks. Associations between prenatal distress measures and maternal faecal bacterial profiles were analysed from 84 maternal faecal specimens collected at the time of delivery. A median of 5465 [interquartile range (IQR): 3159-9877)] post-filtered reads per specimen were obtained following the removal of potential contaminant reads. Infant meconium specimens produced the highest number of post-filtered reads (median: 10002; IQR: 5065–14830), followed by infant faecal specimens collected at 4–12 weeks (median: 6407; IQR: 4069–9042) and 20–28 weeks (median: 5636; IQR: 4228–7084) (35). Maternal faecal specimens sampled at birth produced the lowest number of post-filtered reads (median: 3155; IQR: 2104–4355) (35). All specimen types produced sufficient sequencing depth for calculating Shannon diversity indices in our study (Supplementary Fig. 1).

Demographic and clinical data of the mothers and infants are reported in Table 1. The median (IQR) measures for psychological distress (SRQ-20) and for symptoms of depression (BDI) from mothers with available data (n = 101) were 4.0 (1-7) and 13 (7-12), respectively. Trauma-associated PTSD was diagnosed among 3% (3/101) of mothers. A total of 45% (45/101) of the mothers were exposed to any form of IPV, of which 32% (34/101) experienced emotional abuse, 29% (31/101) physical abuse and 13% (14/101) sexual abuse. Furthermore, 30% (32/101) of the mothers were recently exposed to IPV (during the past 12 months), and 45 % (45/101) had lifetime IPV exposure.

**Table 1:** Demographic and clinical characteristics of mothers and infants included in the study

| Characteristics | Mothers at delivery<br>(n=84) | Infants at birth<br>(n=101) | Infants at 4-12 weeks<br>(n=69) | Infants at 20-28 weeks<br>(n=36) |
| --- | --- | --- | --- | --- |
| Age at which specimens were collected, median (IQR) | 25.3 (21.9 – 31.24)<br>years | 12.4 (4.2 – 24.6) hours | 7.8 (6.9 – 9.3) weeks | 24.1 (23.9 – 26.4) weeks |
| Residential area: |  |  |  |  |
| - Mbekweni, n (%) | 37 (44.0 %) | 54 (53.5 %) | 39 (56.5 %) | 20 (55.6 %) |
| - TC Newman n (%) | 47 (56.0 %) | 47 (46.5 %) | 30 (43.5 %) | 16 (44.4 %) |
| Maternal education: |  |  |  |  |
| - Primary level, n (%) | 8 (9.5 %) | 11 (10.9 %) | 8 (11.6 %) | 7 (19.4 %) |
| - Secondary level, n (%) | 70 (83.4 %) | 82 (81.2 %) | 57 (82.6 %) | 26 (72.2 %) |
| - Tertiary level, n (%) | 6 (7.1 %) | 8 (7.9 %) | 4 (5.8 %) | 3 (8.3 %) |
| Maternal prenatal distress |  |  |  |  |
| -SRQ-20, median (IQR) | 4.0 (2-6) | 4.0 (1-7) | 5 (1-8) | 4.5 (1.3-8) |
| -BDI, median (IQR) | 10 (6-18) | 13 (7-22) | 13 (7-23) | 13 (7-22.5) |
| -PTSD |  |  |  |  |
| -Trauma-exposed, n (%) | 3 (3.6 %) | 3 (3.0 %) | 2 (2.9 %) | 0 (0%) |
| -Suspected PTSD, n (%) | 0 (0 %) | 1 (1.0 %) | 1 (1.4 %) | 1 (2.8 %) |
| -No exposure, n (%) | 81 (96.4 %) | 97 (96.0 %) | 66 (95.7 %) | 35 (97.2 %) |
| -IPV |  |  |  |  |
| -IPV emotional, n (%) | 29 (34.5 %) | 34 (33.7 %) | 24 (34.8 %) | 13 (36.1 %) |
| -IPV physical, n (%) | 26 (31.0 %) | 31 (30.7 %) | 23 (33.3 %) | 10 (27.8%) |
| -IPV sexual, n (%) | 10 (11.9 %) | 14 (13.9 %) | 11 (15.9 %) | 7 (19.4 %) |
| -IPV any, n (%) | 36 (42.9 %) | 45 (44.6 %) | 32 (46.4%) | 18 (50.0 %) |
| -IPV any recent, n (%) | 27 (32.1 %) | 32 (31.7 %) | 25 (36.2 %) | 15 (41.7 %) |
| -IPV any lifetime, n (%) | 36 (42.9 %) | 45 (44.6 %) | 32 (46.4 %) | 18 (50.0 %) |
| Maternal HIV status: |  |  |  |  |
| - HIV infected, n (%) | 18 (21.4 %) | 26 (25.7 %) | 19 (27.5 %) | 10 (27.8 %) |
| Maternal BMI at 6-10 weeks postnatal, median (IQR) | 25.4 (23.2 – 29.2)* | 25.1 (23.0 – 28.8)* | 25.8 (22.9 – 29.7)* | 25.1 (22.4 – 30.4)* |
| Maternal smoking status:# |  |  |  |  |
| - Active smoker (cotinine levels ≥ 500), n (%) | 28 (33.3 %) | 28 (27.7 %) | 21 (30.4 %) | 13 (36.1 %) |
| - Passive smoker (cotinine levels $> 10 < 500$), n (%) | 36 (42.9 %) | 36 (35.6 %) | 19 (27.5 %) | 9 (25.0 %) |
| - Non-smoker (cotinine levels $\leq 10$), n (%) | 20 (23.8 %) | 20 (19.8 %) | 12 (17.4 %) | 6 (16.7 %) |
| Mode of delivery: |  |  |  |  |
| - Vaginal delivery, n (%) | 69 (82.1 %) | 81 (80.2 %) | 57 (82.6 %) | 29 (80.6 %) |
| - Elective caesarean-section delivery, n (%) | 3 (3.6 %) | 5 (5.0 %) | 4 (5.8 %) | 3 (8.3 %) |
| - Emergency caesarean-section delivery, n (%) | 12 (14.3 %) | 15 (14.9 %) | 8 (11.6 %) | 4 (11.1 %) |
| Gestational age (weeks), median (IQR) | 39 (37.3 – 40.0) | 39.0 (38.0 – 40.0) | 39.0 (38.0 – 40.0) | 39.0 (38.3 – 40.0) |
| Birth weight (kg), median (IQR) | 3.0 (2.7 - 3.3) | 3.0 (2.7 - 3.4) | 3.1 (2.7 - 3.5) | 3.0 (2.6 - 3.4) |
| Birth length (cm), median (IQR) | 50.5 (48.0- 52.8) | 50 (48.0 – 53.0) | 51 (48.5 – 53.0) | 50.5 (49 - 53) |
| Sex: |  |  |  |  |
| - Male, n (%) | 36 (42.9 %) | 43 (42.6 %) | 30 (43.5 %) | 18 (50 %) |
| Milk feeding: |  |  |  |  |
| - Exclusive breastfeeding, n (%) | 71 (84.5 %)§‡ | 81 (80.2 %)§‡ | 32 (46.4 %)§φ | 6 (16.7 %)§† |
| - Exclusive formula feeding, n (%) | 9 (10.7 %)§‡ | 16 (15.8 %)§‡ | 11 (15.9 %)§φ | 7 (19.4 %)§† |
| - Mixed feeding□, n (%) | - | - | 25 (36.2 %)§φ | 23 (63.9 %)§† |
| Antibiotic use, n (%) | 9 (10.7 %) | 15 (14.9 %) | 9 (13.0 %) | 3 (8.3 %) |
| Household members, median (IQR) | 5 (3 - 7) | 4 (3 - 6) | 4 (3 - 6) | 4 (3 - 6) |

### Association of maternal prenatal psychological measures with infant faecal bacterial profiles

Infants born to mothers with exposure to lifetime IPV had higher proportions of unclassified genera within the family Enterobacteriaceae [OTU 101 (p < 0.01); OTU 615 (p = 0.02); OTU 616 (p = 0.01)] measured from meconium (Fig. 1 A-C) (covariates significantly associated with prenatal psychological measures or bacterial taxa that were included in the statistical model: maternal BMI and breastfeeding).

**Figure 1.**
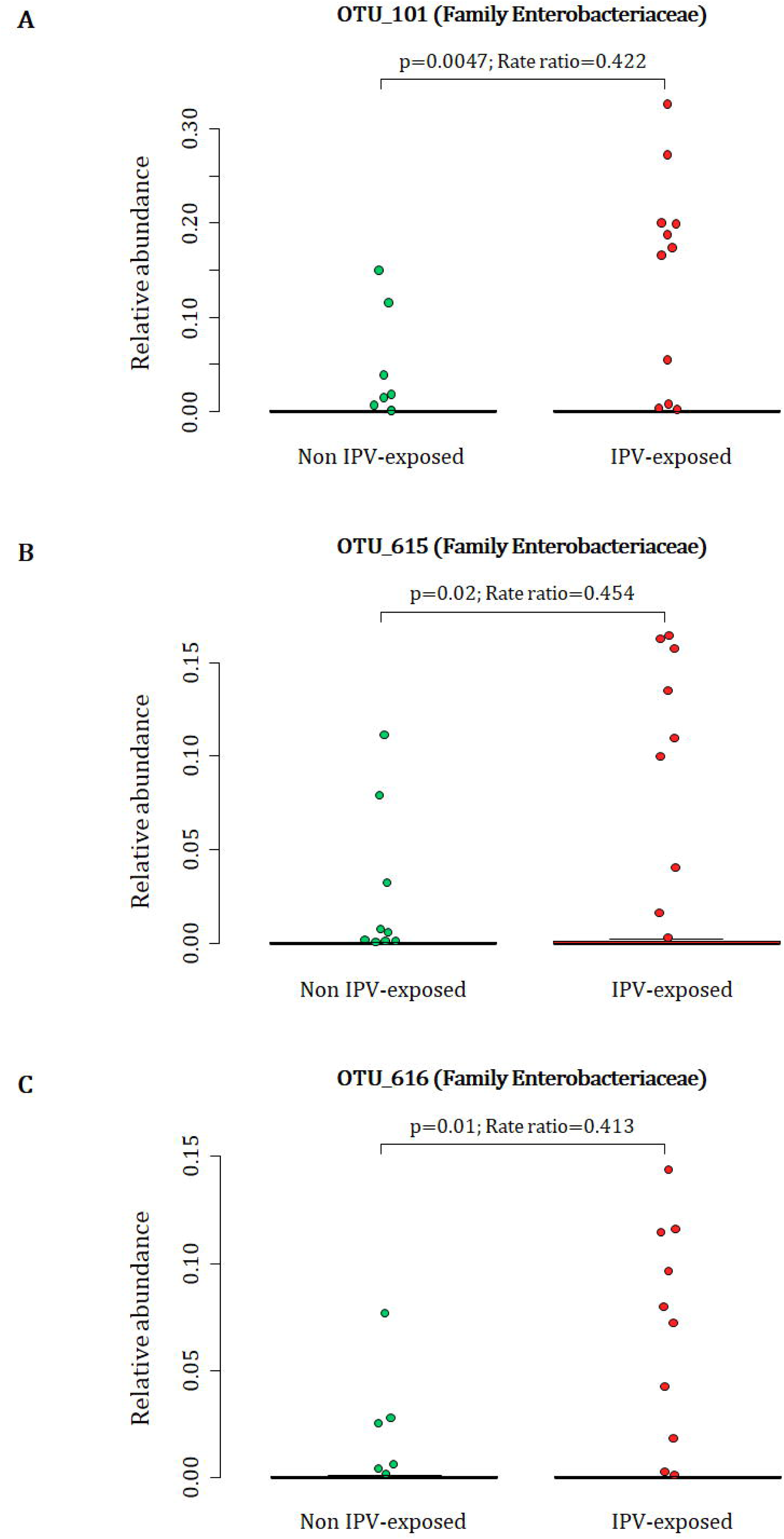
Relationship between maternal lifetime exposure to IPV and the infant meconium bacterial profiles. The number of participants represented in this figure were infants from which we detected the respective OTUs at proportions > 0. A) Differences in proportions of OTU 101 detected from 88 infant meconium specimens; B) differences in proportions of OTU_615 detected from 73 infant meconium specimens; and C) differences in proportions of OTU_616 detected from 71 infant meconium specimens at proportions > 0. IPV=intimate partner violence.

A total of 88 participants at birth had OTU 101, 73 infants had OTU 615 and 71 infants had OTU 616 detected at proportions > 0 from their meconium (Fig. 1 A-C). At 4-12 weeks of age, infants born to mothers with exposure to lifetime IPV had higher proportions of the genus *Weissella* (Phylum Firmicutes; Class Bacilli; Order Lactobacillales; Family Leuconostocaceae) (p < 0.001) when compared to infants born to mothers who were not exposed to lifetime IPV (Fig. 2) (covariates significantly associated with prenatal psychological measures, bacterial taxa or diversity that were included in the statistical model: area and maternal HIV status). A total of 44 infants at 4-12 weeks of age had *Weissella* detected at proportions > 0 from their faecal specimens (Fig. 2). This association was not found to be significant at birth or at 20-28 weeks (Supplementary figure 2).

**Figure 2.**
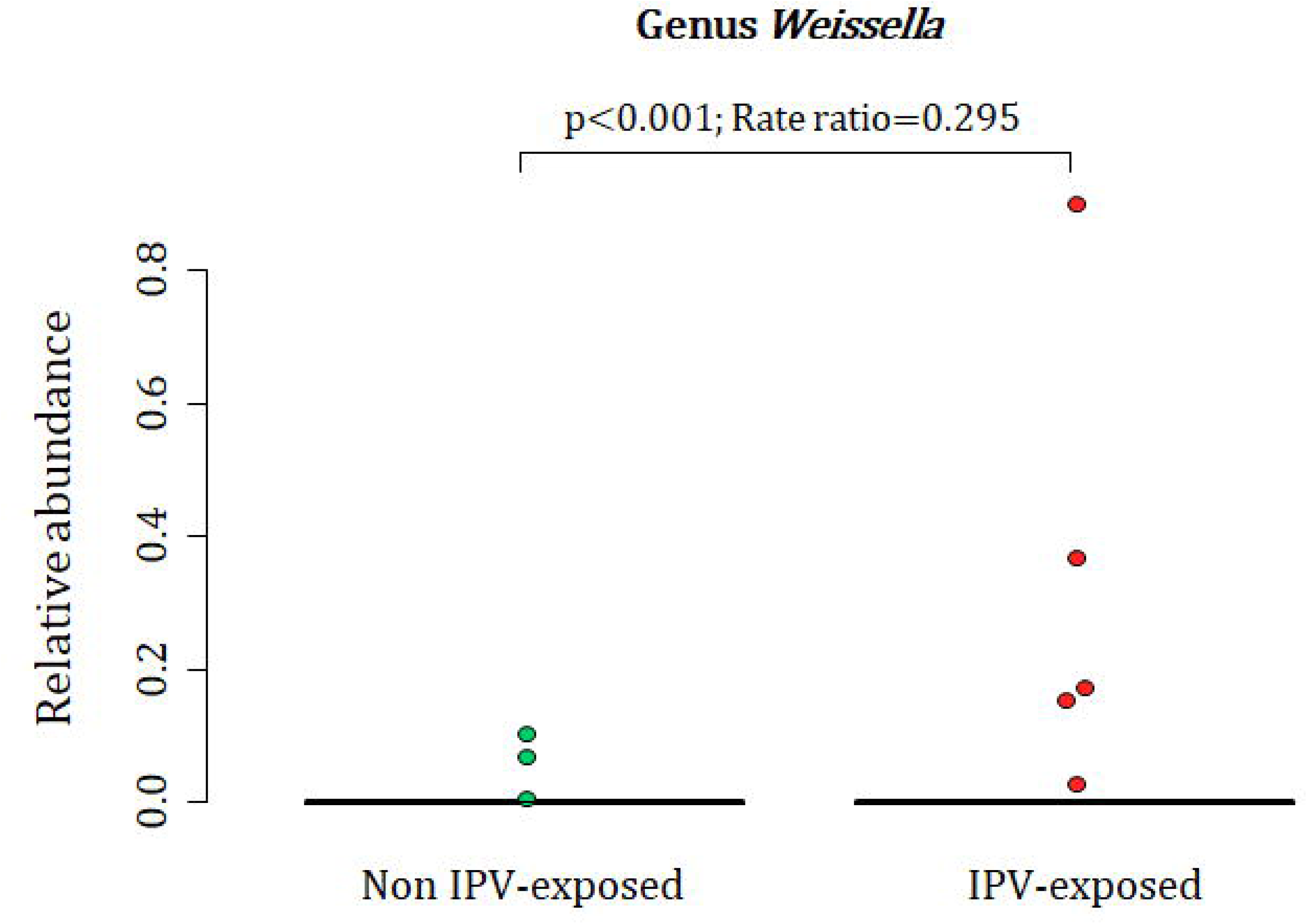
Relationship between maternal lifetime exposure to IPV and the infant faecal bacteria at 4-12 weeks. The figure presents differences in proportions of genus *Weissella* that was detected from 44 infant faecal specimens at proportions > 0. IPV=intimate partner violence.

At 20-28 weeks, increased measures of maternal psychological distress (SRQ-20) were significantly associated with lower proportions of the family Veillonellaceae (Phylum Firmicutes, Class Negativicutes (p = 0.003), Order Selenomondales (p = 0.003)) (p = 0.01) (covariates significantly associated with psychological measures, bacterial taxa or diversity which were included in the statistical model: area, maternal HIV status, maternal education and gender of infant) (Fig. 3 A-C). This association was only significant at 20-28 weeks (Supplementary figure 3).

**Figure 3.**
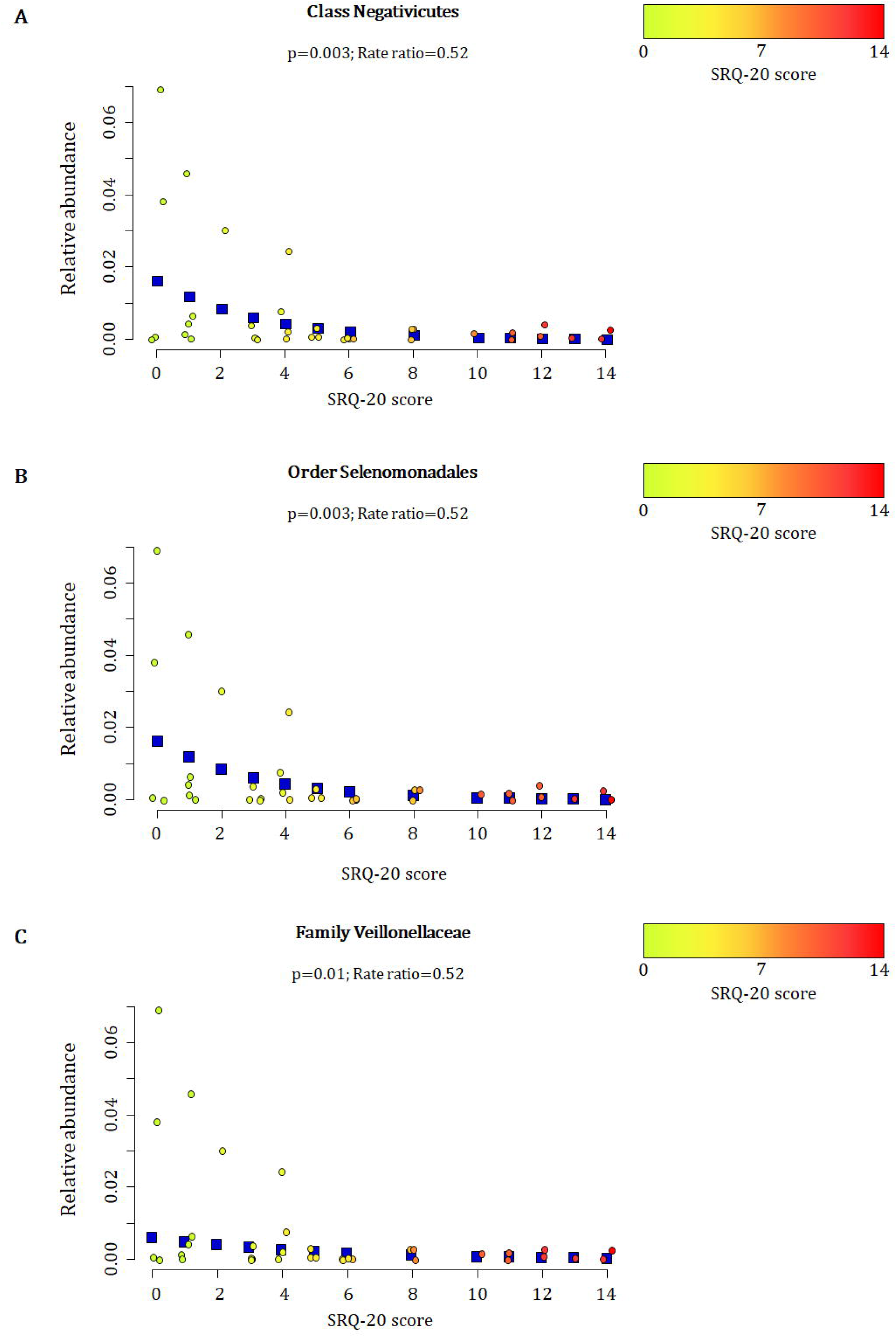
Relationship between maternal prenatal psychological distress (SRQ-20) and infant faecal bacterial profiles at 20-28 weeks. Proportions of the A) class Negativicutes, B) order Selenomonadales, and C) family Veillonellaceae observed from faecal specimens collected at 20-28 weeks were inversely correlated with maternal psychological distress (SRQ-20) (p ≤ 0.01). Individual maternal SRQ-20 scores for the 36 infants investigated at 20-28 weeks are represented by circles using a gradient of colours ranging from green (low SRQ-20 scores) to red (high SRQ-20 scores). Blue squares represent the non-parametric regression smoother used to show the trend in the observations.

When measuring the effect of maternal prenatal stressors and distress on n=36 infant faecal bacterial profiles over time, we found that maternal prenatal exposure to lifetime IPV was associated with longitudinal increases of taxa within the family Enterobacteriaceae (genus Citrobacter (p = 0.002) (Fig. 4 A) and unclassified genera OTU 101 (p = 0.006); OTU 615 (p=0.003); OTU 616 (p = 0.003)) (covariates significantly associated with psychological measures or bacterial taxa that were included in the statistical model: gender, area and maternal HIV status) (Fig.4 B-D) in infants over time. Infants from mothers that were exposed to lifetime IPV had higher portions of the family Staphylococcaceae at birth compared to proportions at ages 4-12 weeks (p=0.038) and 20-28 weeks (p=0.018) (Fig. 4 E). Higher scores of maternal psychological distress were associated with a slower decline of Gammaproteobacteria between birth and 4-12 weeks (p = 0.007) and between birth and 20-28 weeks (p = 0.089) (covariates significantly associated with psychological measures or bacterial taxa that were included in the statistical model: gender, area and maternal HIV status) (Fig 4 F).

**Figure 4.**
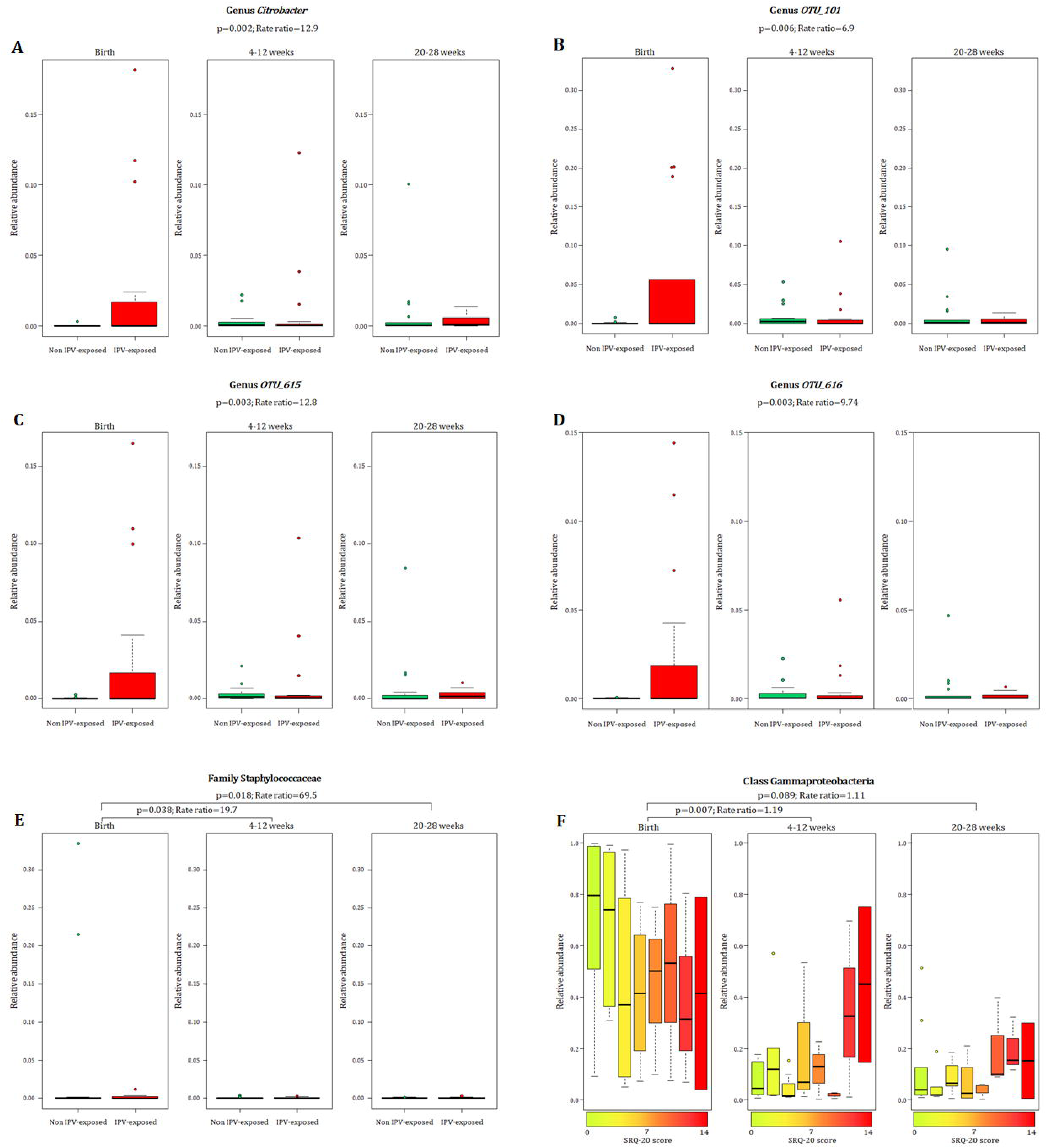
Relationship between maternal exposure to lifetime IPV, psychological distress and temporal changes in infant faecal bacterial profiles. Longitudinal changes to the proportions of A) Genus *Citrobacter* and unclassified genera within the family Enterobacteriaceae B-D) [OTU_101 (p = 0.006); OTU_615 (p=0.003); OTU_616 (p = 0.003)] in the infants born to mothers exposed to lifetime IPV. E) Proportions of the family Staphylococcaceae at birth compared to proportions at ages 4-12 weeks (p=0.038) and 20-28 weeks (p=0.018) of infants from mothers exposed to lifetime IPV. F) Association of maternal prenatal psychological distress (SRQ-20) with faecal Gammaproteobacteria in infants over time. Individual maternal SRQ-20 scores for the 36 infants are represented by using a gradient of colours ranging from green (low SRQ-20 scores) to red (high SRQ-20 scores). IPV = intimate partner violence.

No significant associations with infant faecal bacterial profiles were found when assessing symptoms of depression or PTSD. Supplementary figures 4-8 show different measures for maternal prenatal stressors and distress with the infant microbiota at genus level measured over time. No significant associations were found between any maternal prenatal psychological measures and infant faecal bacterial diversity indices (data not shown).

### Prenatal psychological measures and maternal faecal bacterial profiles at delivery

Maternal exposure to recent IPV (past year) was associated with higher proportions of the family Lactobacillaceae (p = 0.03) (phylum Firmicutes, class Bacilli, order Lactobacillales) and lower proportions of the family Peptostreptococcaceae (p = 0.04) (phylum Firmicutes, class Clostridia, order Clostridiales) in maternal faecal specimens at delivery (covariates significantly associated with psychological measures or bacterial taxa that were included in the statistical model: mode of delivery) (Fig 5. A and B). None of the prenatal psychological measures were associated with maternal faecal bacterial diversity indices (data not shown).

**Figure 5.**
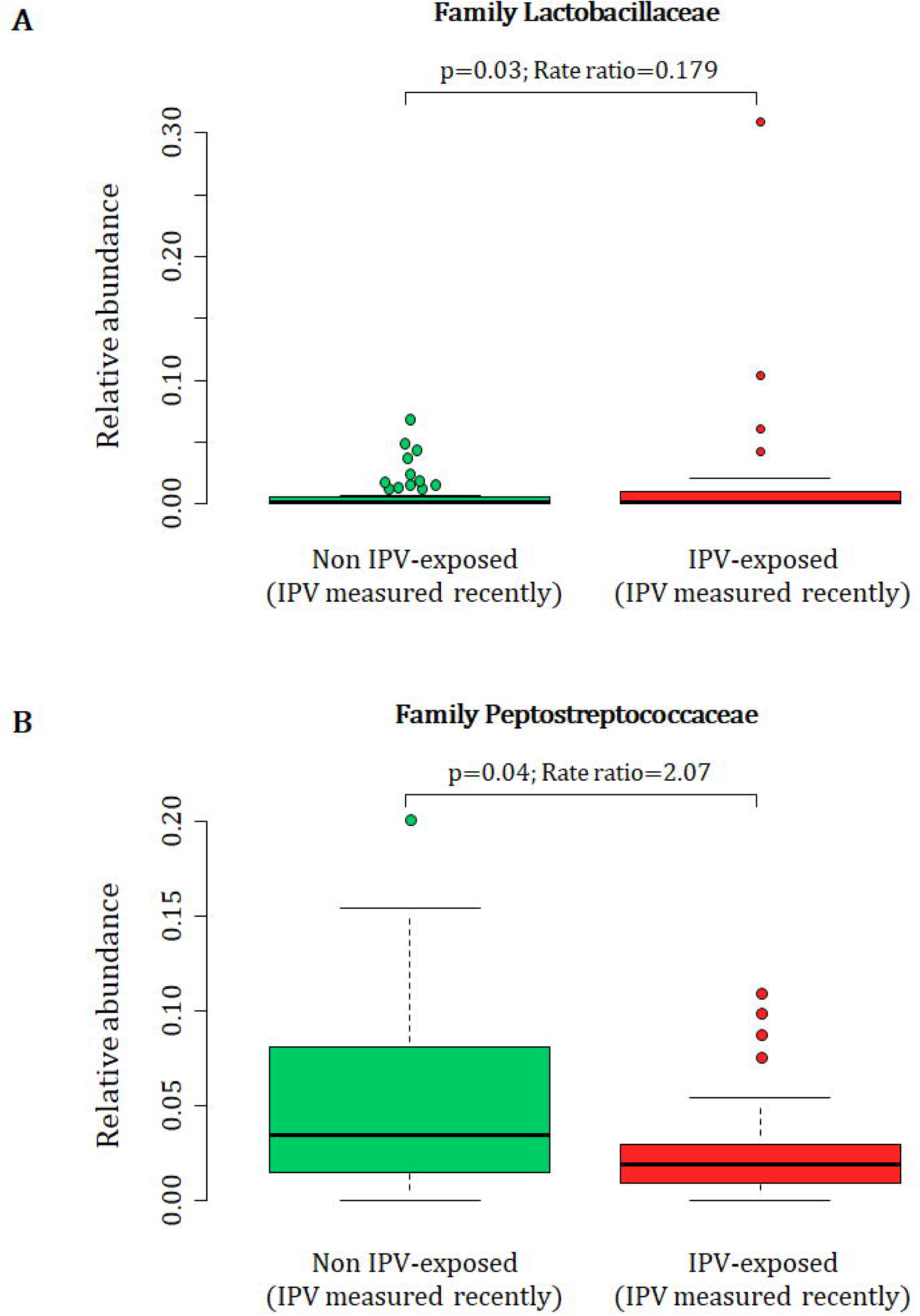
Relationship between maternal exposure to recent IPV (past year) during pregnancy and maternal faecal bacterial profiles at delivery. IPV = intimate partner violence.

### Beta diversity and measures for maternal psychological adversities

No clustering patterns were apparent for maternal or infant faecal bacterial profiles in relation to any of the prenatal measures for psychological adversities based on the “w’ metric for beta diversities (results not shown).

## Discussion

This study characterized the association of maternal prenatal stressors and distress with early life infant faecal and maternal bacterial profiles. Our results demonstrate that maternal exposure to lifetime IPV and psychological distress were significantly associated with early infant faecal bacterial profiles.

Until recently, the foetus has been considered to exist in a sterile environment with initial microbial colonization of the newborn taking place during birth (51). Recent studies have reported the detection of microbes from amniotic fluid, placenta and meconium, suggesting that microbial colonization may begin in utero (52, 53). In addition, a number of studies have reported bacterial communities from meconium specimens (54). Dysbiosis of faecal bacterial profiles early in life has been associated with a number of neonatal diseases, including asthma, allergic diseases, and obesity (51), which further emphasises the importance of *in utero* colonization on the health of newborns.

Our findings in relation to the family Enterobacteriaceae are of particular interest, given previous findings. We detected higher abundances of unclassified taxa (OTU 101, OTU 615 and OTU 616) within the family Enterobacteriaceae (within the phylum Proteobacteria) from meconium specimens of infants born to mothers exposed to IPV. We further showed that the same taxa, as well as the genus Citrobacter (also a member of the Enterobacteriaceae) increased significantly over time amongst infants born to mothers experiencing IPV. A higher cumulative score of self-reported maternal stress during pregnancy has previously been associated with increased relative abundances of Proteobacteria in infant faecal specimens over time (10). The family Enterobacteriaceae (within the phylum Proteobacteria) has also been associated with depression in an adult population (55), and leakage of lipopolysaccharide from Enterobacteriaceae into the circulation has been associated with major depression. Increased faecal Enterobacteriaceae in infants at 3 to 6 months of age has also been associated with impaired follow-up fine motor skills in the children at 3 years (14). Our findings on the other hand differ from a study by Hu and colleagues that showed an association of higher scores for pregnancy-related anxiety with lower abundances of an unclassified genus in the family Enterobacteriaceae from meconium of 75 infants (11). This discrepancy may be due differences of the psychological measures and possibly the larger proportion of mothers (27% vs. 11%) that used antibiotics prior to delivery compared to our study.

We also found that infants born to mothers with IPV exposure had higher proportions of the genus *Weissella* (phylum Firmicutes) at 4-12 weeks of age. The relative abundance of *Weissella* increased in all infants with age. However, this increase occurred earlier for infants born to mothers with IPV exposure and subsequently reached the same level for all infants at 20-28 weeks of age. To the best of our knowledge, no prior associations have been reported between psychological stressors or distress and the genus *Weissella*. Furthermore, premature delivery has been associated with distinct infant faecal bacterial profiles, including higher abundances of *Weissella* (56). However, the infants assessed in this study sample had a median gestational age of 39 weeks (IQR: 38-40) (35). Moreover, there were no significant age differences (p > 0.5) between infants from mothers that were IPV-exposed vs. non IPV-exposed at any of the time points at which specimens were collected.

Our results further show that stool from infants born to mothers with higher measures of psychological distress (SRQ-20) showed a slower decline in Gammaproteobacteria over time compared to infants born to mothers with lower SRQ-20 levels. The class Gammaproteobacteria (and family Enterobacteriaceae in particular) are the most abundant faecal bacteria in the first month of life, subsequently substituted by anaerobes (57). Slower reduction rates of these bacteria over time may be an indication of impaired maturation of the infant’s gut bacteriome (57). Furthermore, higher measures of maternal prenatal psychological distress were associated with lower relative abundance of Veillonellaceae (phylum Firmicutes) in infant faecal specimens at 20-28 weeks. The median relative abundance of Veillonellaceae increased with age overall amongst infants in this study. However, at the age of 4-12 weeks, this abundance was relatively lower (not significant) in infants from mothers with higher scores for psychological distress and more pronounced and significantly lower in these infants at the age of 20-28 weeks. Veillonellaceae are lactate utilizing bacteria that are abundant in infant faecal specimens up to 6 months and subsequently decrease during the transition to toddler and then remaining at stable levels throughout healthy adulthood (58-60). Decreased abundances of Veillonellaceae have been found in children with autism (61), as well as in patients with depression (55). A recent Belgian population cohort showed that the genus *Dialister* (within the family Veillonellaceae) was significantly deceased in depression, a finding that was further validated in a large Dutch cohort (62). These findings suggest a consistent association between decreased abundance of Veillonellaceae and neurodevelopmental impairment or psychological distress.

Overall, our results illustrate that the associations of different psychological profiles during pregnancy on faecal bacteria may vary. Exposure to lifetime IPV in particular, was associated with differences in the infant and maternal faecal bacteria. IPV is a traumatic stressor that often occurs over prolonged and repeated episodes (63) and IPV during pregnancy often persists postpartum (64-66). It should be noted that other effects of IPV which were not controlled for in this study may contribute to its association with changes to infant faecal bacterial composition. For example, mothers who have experienced IPV are more likely to have weakened bonding with their infant (66, 67), which may in turn affect the infant faecal bacterial composition. Supporting evidence from animal work showed that male rat pups that were stressed by maternal separation had elevated faecal levels of Enterobacteria and Bacteroides during adulthood (68). Another animal study found that maternal separation of pups resulted in a reduced ratio of Firmicutes to Bacteroides in the adult gut of these female rats (69). The effect of IPV on child feeding may also contribute to faecal bacterial changes. Studies have showed that maternal IPV can contribute to infant malnutrition in low to middle income countries (66, 70, 71). Malnutrition has been associated with marked changes in infants’ faecal bacterial composition (72). Therefore, other measures that accompany IPV post-partum may play a role in the development of infant faecal bacterial composition.

Several limitations to our study deserve emphasis. First, there were missing faecal specimens over time, due mainly to the inability to collect stool at the scheduled study visits (stool was only collected if the infant was able to produce a stool sample during the study visit). Therefore, our results should be interpreted with caution, especially at the 20-28 weeks interval. Second, given the limited number of mothers with PTSD, our analysis has insufficient statistical power to draw definitive conclusions about the possible association between PTSD and an altered faecal bacteriome. Third, the cross-sectional analysis for the individual time points does not allow for definitive conclusions about their longitudinal changes in the faecal bacterial profiles. Further work is needed to understand whether IPV is a proxy for other key variables that may affect the associations found with the infants’ faecal gut bacteria.

In summary, maternal prenatal psychological stressors and distress, particularly IPV during pregnancy, is associated with early changes in infant faecal bacterial profiles in a semi-rural South African area. Further studies are needed to verify our findings, and to determine wheteher these changes to the gut bacteriome due to maternal prenatal psychological stressors and distress are associated with neurodevelopmental impairments in children over time. In addition, it would be of interest to examine if preventative strategies to reduce the risk of prenatal psychological stressors and distress can prevent longitudinal changes to the infant’s gut bacteria and associated health risks. Such work may provide further insights into understanding relevant mechanisms and identifying appropriate targets, with the ultimate aim of preventing the negative effects of intergenerational transmission of psychopathology on the developing brain.

## Acknowledgements

We would like to thank the families and their children that participated in this study. We thank the study staff, administrative staff and clinical staff of the Western Cape Government Health Department at TC Newman and Mbekweni clinics, and Paarl Hospital for their support of the study. Petrus Naudé, Shantelle Claassen-Weitz, Mamadou Kaba, Heather Zar, Mark Nicol and Dan Stein contributed to the concept and design of the study. Shantelle Claassen-Weitz, Sugnet Gardner-Lubbe, Gerrit Botha and Mamadou Kaba contributed to the acquisition and analysis of data. All mentioned authors contributed to the interpretation of the results presented in this manuscript. Petrus Naudé and Shantelle Claassen-Weitz drafted the manuscript. The critical revisions by Sugnet Gardner-Lubbe, Gerrit Botha, Mamadou Kaba, Heather Zar, Mark Nicol and Dan Stein were essential for the intellectual content of this manuscript. All listed authors provided final approval of the version to be published.

## Ethical standards

The authors assert that all procedures contributing to this work comply with the ethical standards of the relevant national and institutional committees on human experimentation and with the Helsinki Declaration of 1975, as revised in 2008.

