## Supplementary material for "Association of maternal prenatal psychological stressors and distress with maternal and early infant faecal bacterial profile"

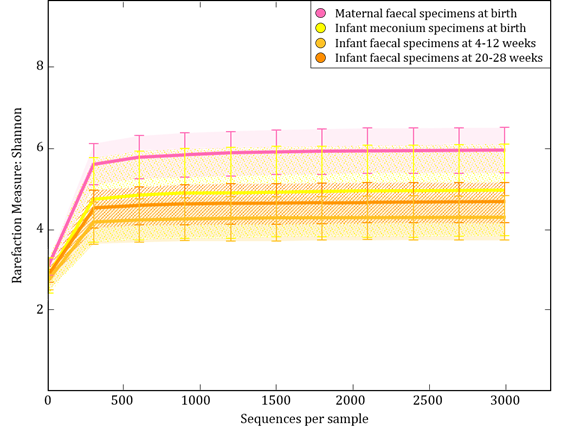

Supplementary figure 1. Rarefaction curve analysis indicates sufficient sequences for calculating Shannon diversity indices. Alpha diversity indices were previously compared between the different groups as depicted in the figure (1). Maternal faecal specimens had significantly higher alpha diversity indices compared to infant meconium specimens, while infant meconium specimens had significantly higher alpha diversities compared to infant faecal specimens collected at 4-12 and 20-28 weeks of life (1).

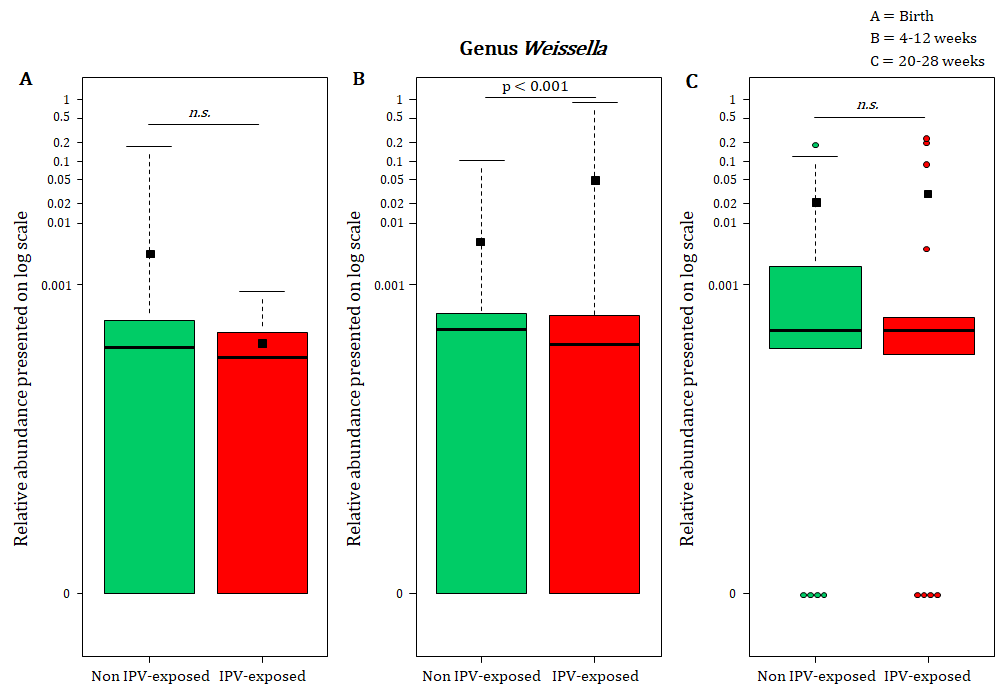

Supplementary figure 2. Maternal lifetime exposure to IPV and *Weissella* in the infant faecal bacteria at (A) birth, (B) 4-12 weeks and at (C) 20-28 weeks. Values are presented on a log scale. IPV; intimate partner violence, *n.s.*; not significant.

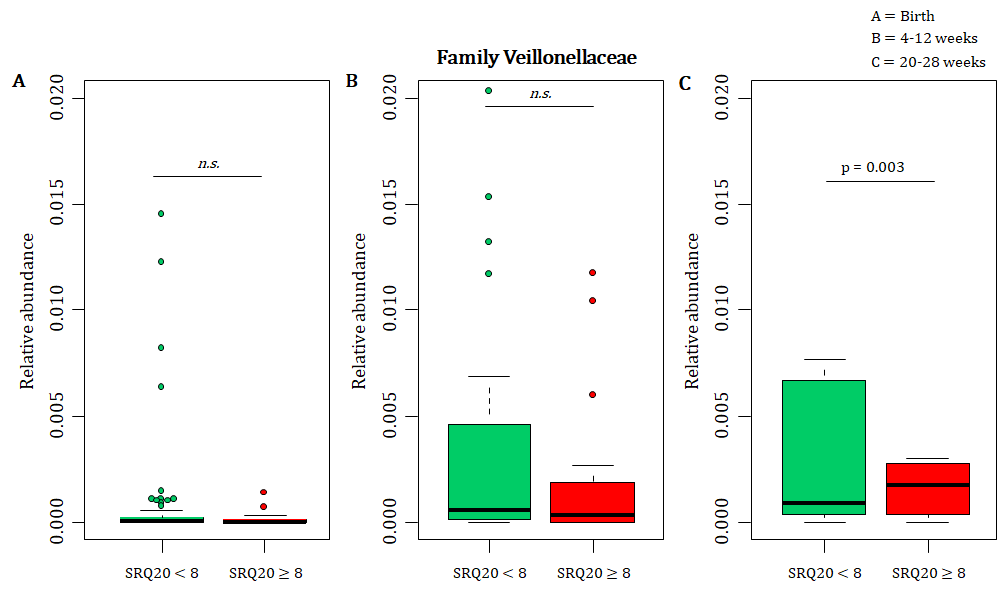

Supplementary figure 3. Relationship between maternal prenatal psychological distress (SRQ-20) and abundances of infant faecal Veillonellaceae at (A) birth, (B) 4-12 weeks and at (C) 20-28 weeks. *n.s.*; not significant.

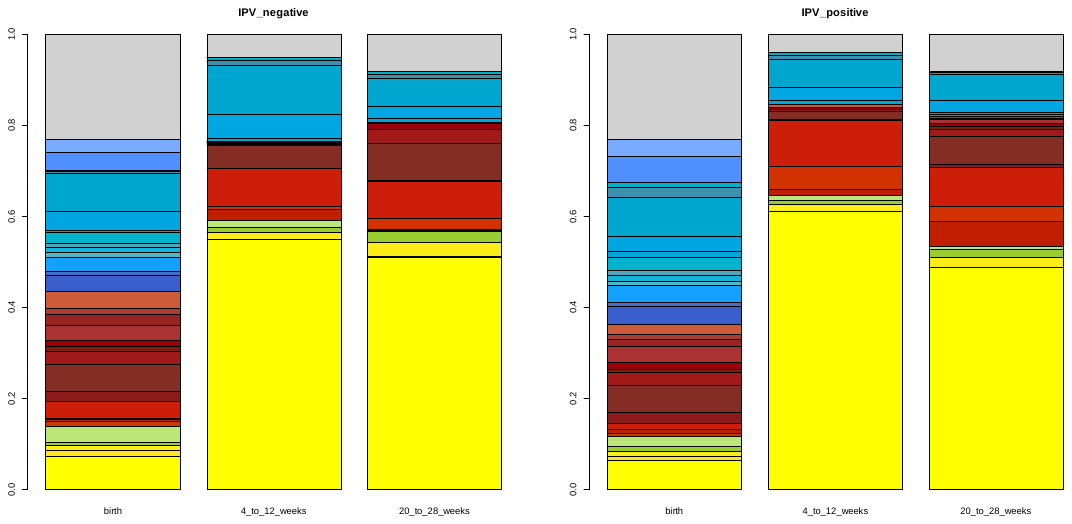

B

A

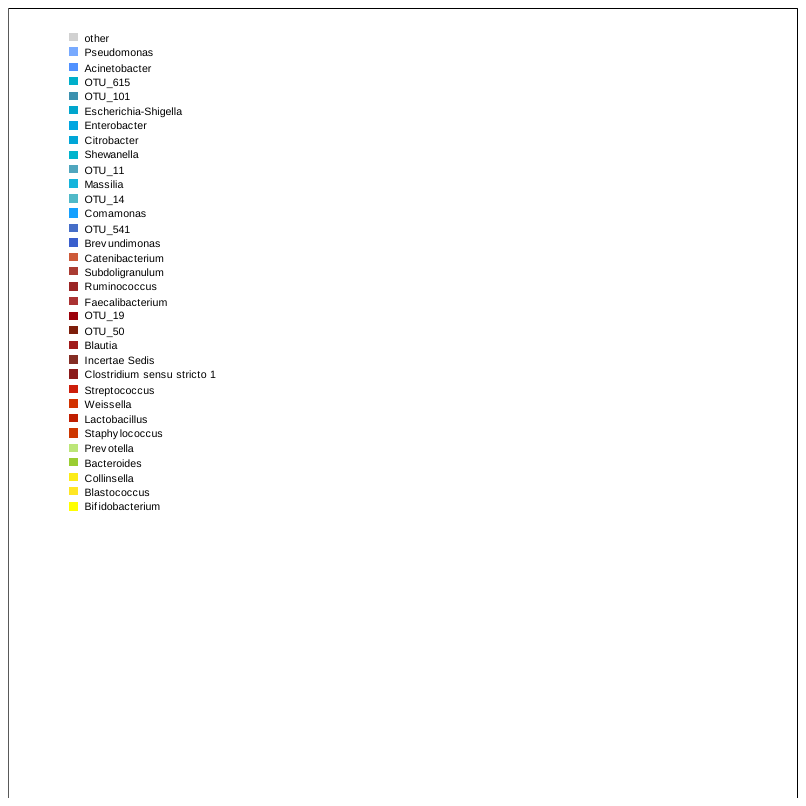

Legend:

Supplementary figure 4. Average relative abundances of genus-level faecal bacteria in 36 infants with longitudinal data collected from mothers with A) no/low lifetime exposure to intimate partner violence (IPV) (n=18) vs. B) high lifetime exposure to IPV (n=18).

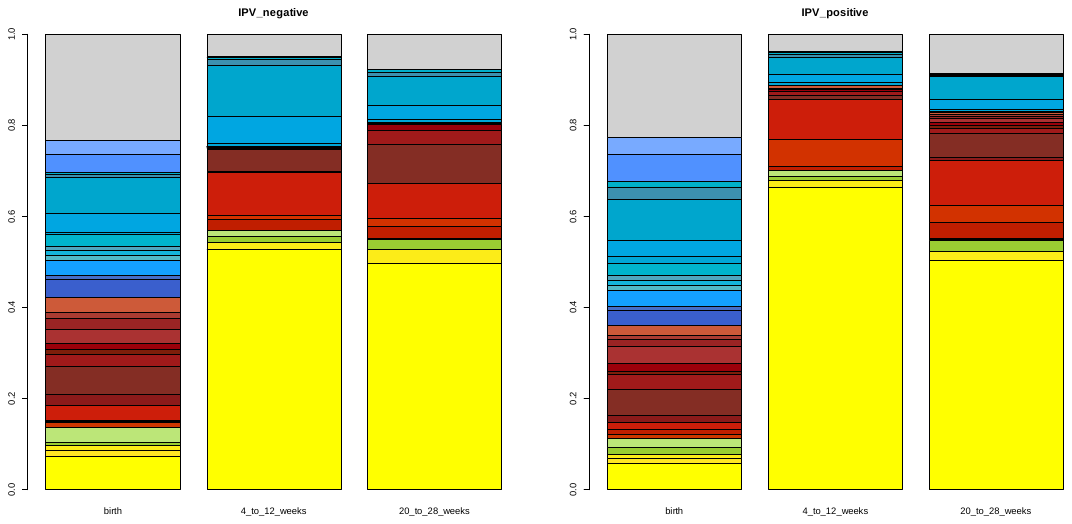
Supplementary figure 5. Average relative abundances of genus-level faecal bacteria in 36 infants with longitudinal data collected from mothers with A) no/low recent intimate partner violence (IPV) (past year) exposure (n=21) vs. B) high recent IPV (past year) exposure (n=15).

A

B

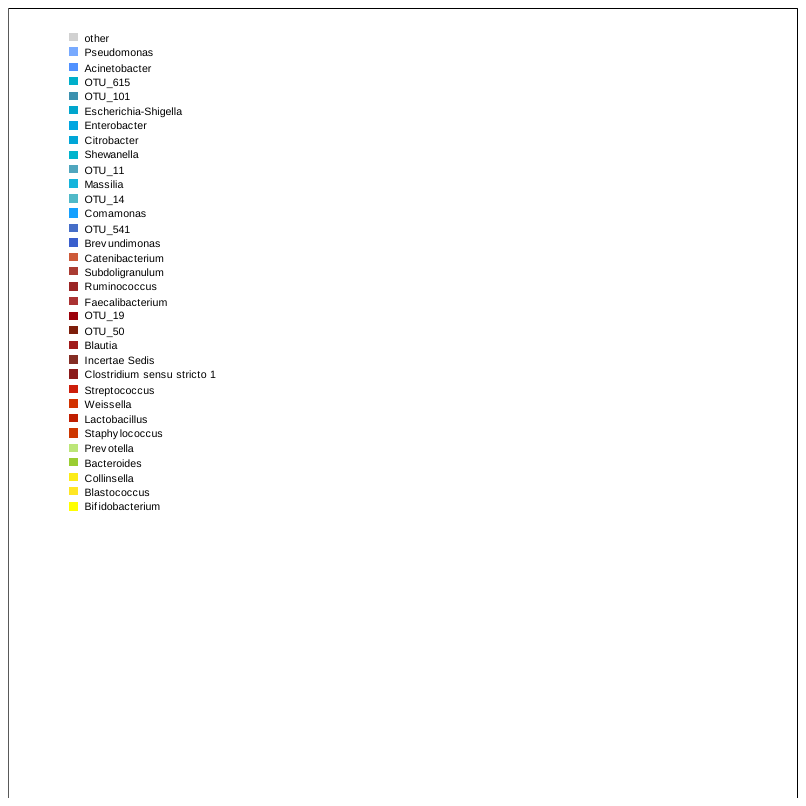

Legend:

Supplementary figure 6. Average relative abundances of genus-level faecal bacteria in 36 infants with longitudinal data collected from mothers with A) no post-traumatic stress disorder (PTSD) (n=11) vs. B) PTSD (n=25).
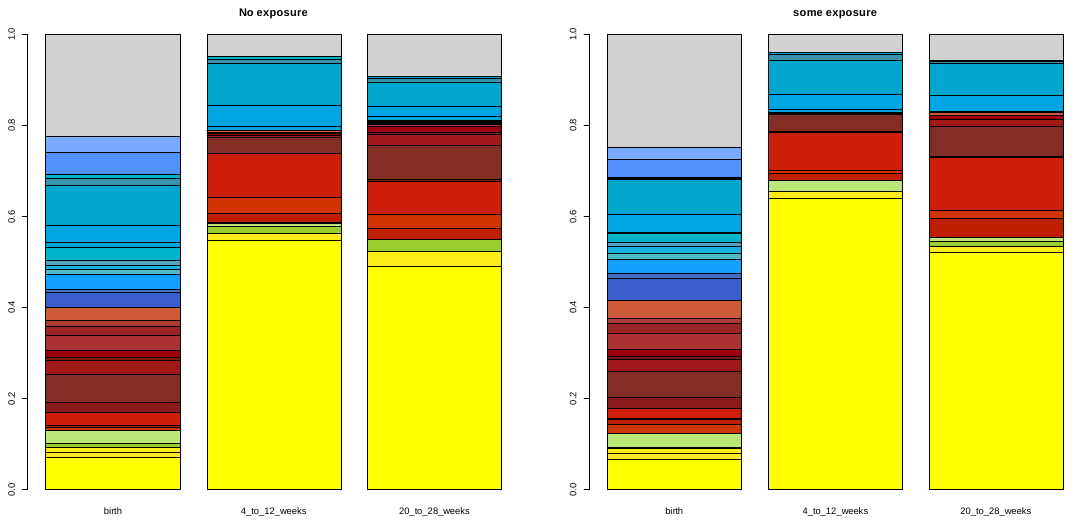
 PTSD were dichotomized according to non-exposed vs. trauma-exposed and suspected PTSD.

B

A

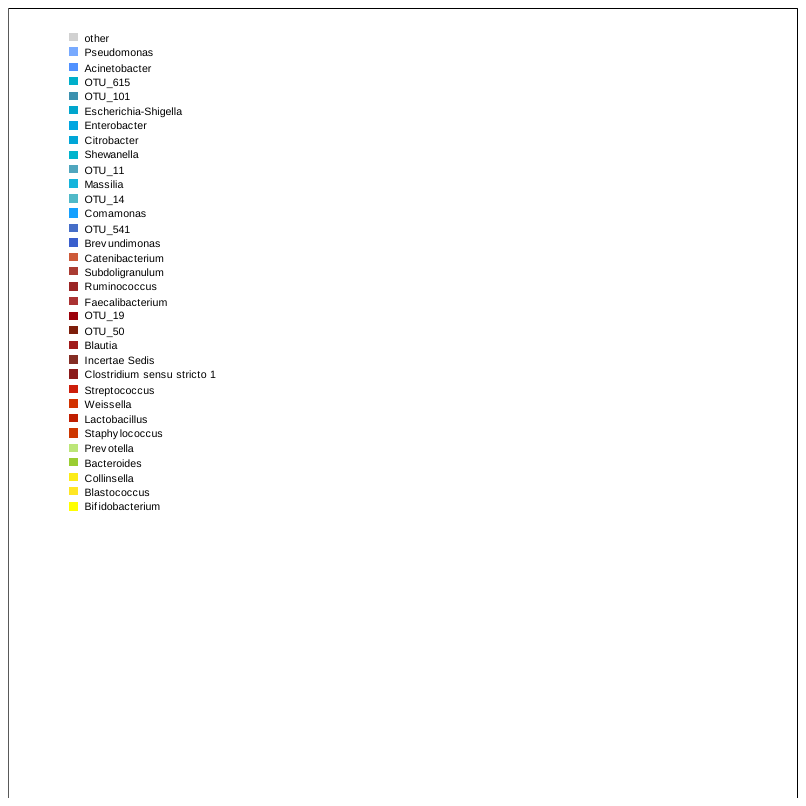

Legend:

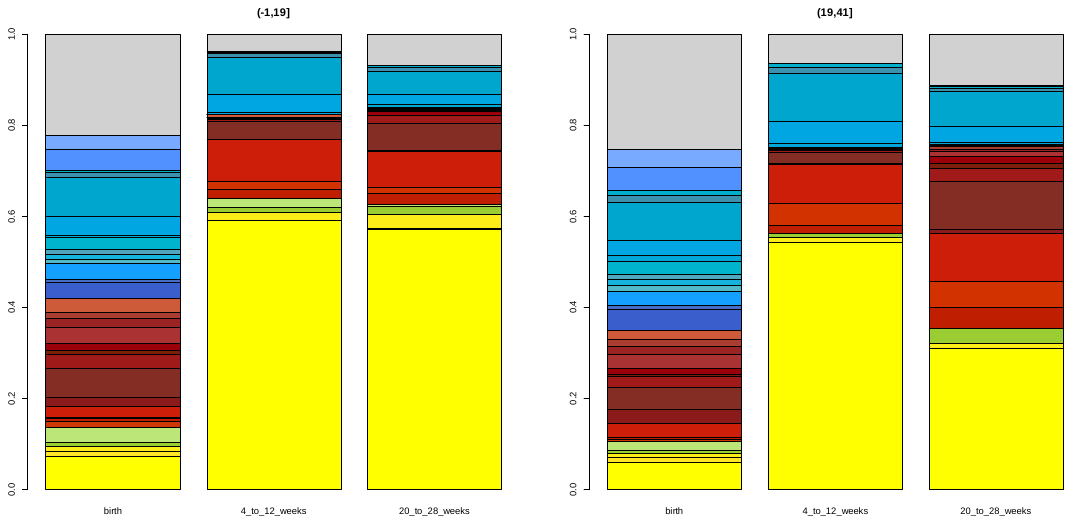
Supplementary figure 7. Average relative abundances of genus-level faecal bacteria in 36 infants with longitudinal data collected from mothers with A) no symptoms of depression (BDI) (n=26) and B) symptoms of depression (n=10). A cut-off score of ≥20 was used to dichotomize participants into “probable moderate/severe clinical cases” versus “probable sub-threshold participants”

B

A

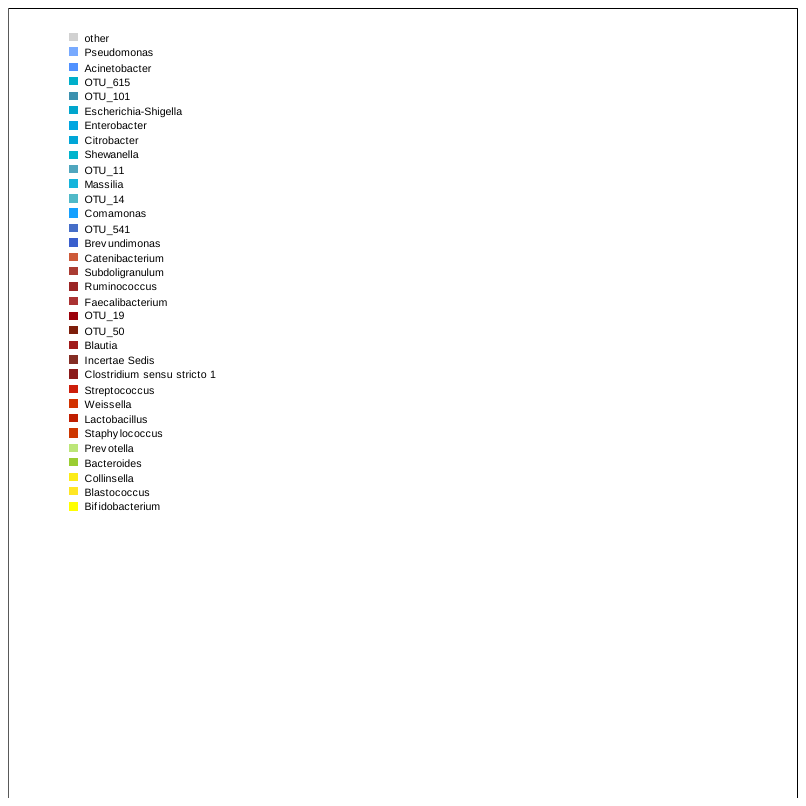

Legend:

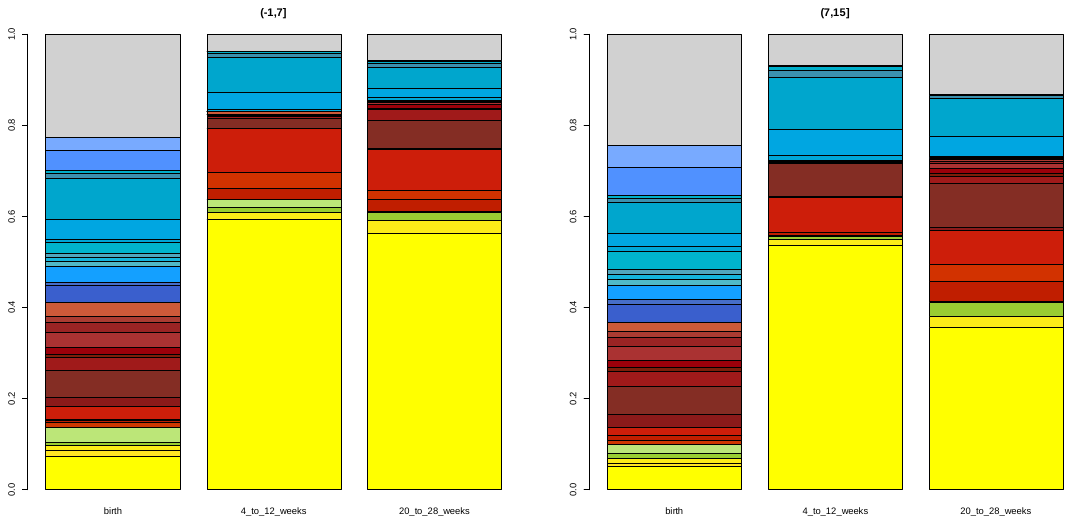
Supplementary figure 8. Average relative abundances of genus-level faecal bacteria in 36 infants with longitudinal data collected from mothers with A) low risk for psychological distress (SRQ-20) (n=25) and B) with high risk for psychological distress (n=11). An SRQ-20 cut-off score of <8 was used to dichotomize participants into “low risk” versus “high risk”.

A

B

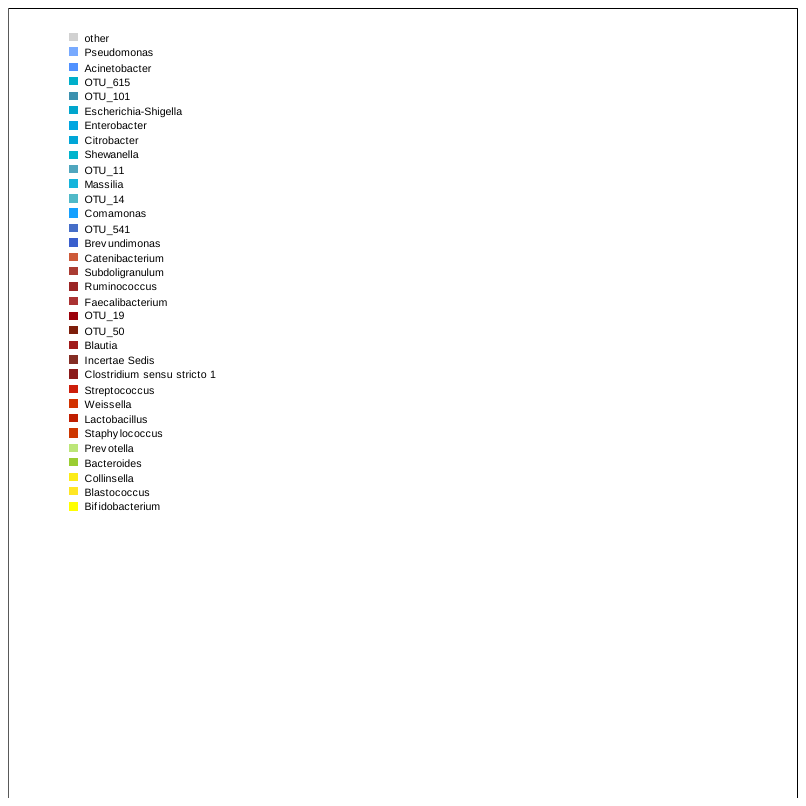

Legend:
